# Genetic Mapping of All Human Paralog Pairs to Characterize Synthetic Lethality and Buffering

**DOI:** 10.1101/2024.07.16.603642

**Authors:** Michael J. Flister, Daniel Verduzco, Sakina Petiwala, Tatiana C. Carneiro-Lobo, Christos Ghekas, Xu Shi, Charles Lu, Zoltan Dezso

## Abstract

Paralogs are abundant in the human genome and thought to be a primary source of synthetic lethality, yet the vast paralogome remains largely uncharacterized. A digenic screen of all paralogous gene pairs in the human genome revealed that synthetic lethalities were infrequent and varied in penetrance in different tumor backgrounds. We hypothesized that the variable penetrance of synthetic lethalities resulted from complex polygenic interactions with different cellular contexts. A machine learning classifier of a subset of paralog pairs tested across 49 cancer models revealed that endogenous perturbations in related pathways predicted paralog synthetic lethality. Further, predictive modeling of paralog synthetic lethality revealed that the strength of synthetic lethal interactions were largely due to the overlap and essentiality of the protein-protein interaction networks shared by the paralogs pairs. Collectively, this study tested all digenic paralog interactions and delineated the key feature classes that underlie the heterogeneity of paralog synthetic lethalities.

**Highlights:**

- Synthetic lethal paralogs have long been known, yet most paralog interactions are untested.
- We report the first combinatorial screen of all 36,648 known human paralog pairs.
- A meta-analysis of 461 paralog pairs in 49 cell models revealed context-dependent interactions.
- The essentiality of paralog interaction networks dictate the strength of synthetic lethality.

**eTOC:** Synthetic lethal paralogs have long been known, yet most paralog interactions are untested. Flister et al report a screen of all 36,648 known human paralog pairs. These data combined with a meta-analysis of 461 pairs across 49 cell models revealed insights to the molecular underpinnings of context-dependent synthetic lethality.

**Graphical Abstract:** 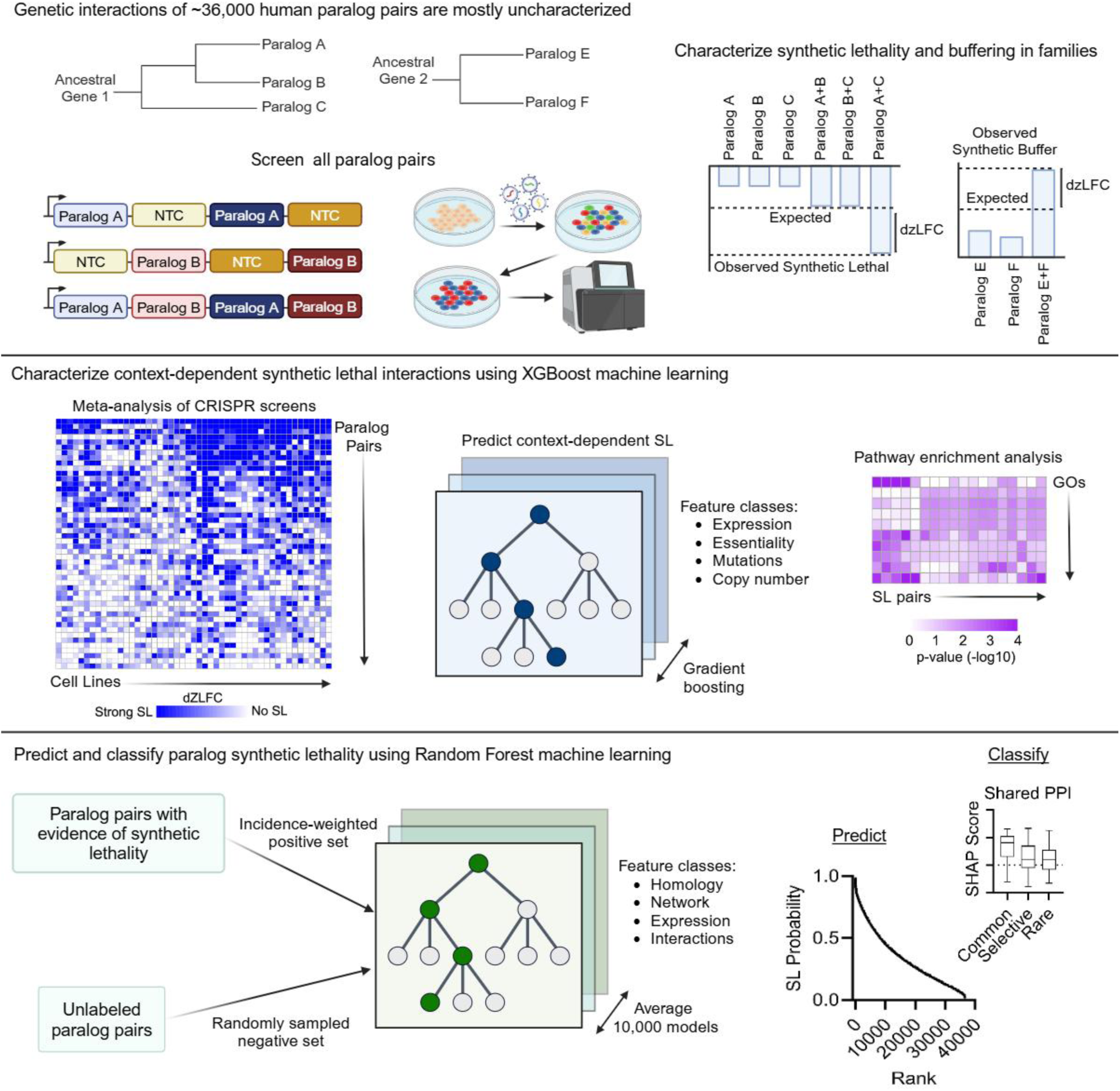

## INTRODUCTION

Paralogs are functionally related genes that arise from genomic duplications and serve as a major source of genetic evolution through selective adaptations in most species, including humans^1,2^. The human genome has an estimated 13,320 paralogs belonging to ∼3,500 families (ranging from 2 to 48 members)^3^. Many parlogs are thought to share compensatory functions and are therefore more likely to exhibit synthetic lethality (i.e., the loss of one paralog conveys dependency to another)^4^. The loss of a paralog in tumor cells can be driven by a selective advantage or as a genomic passenger that is co-deleted with another tumor suppressor gene, which renders the tumor cells sensitive to selective inhibition of the remaining paralog. Because normal cells retain both paralogs, a selective inhibitor of a single paralog is well tolerated in normal tissues, whereas it is synthetic lethal in the tumor cells^2,5^.

Although the concept of synthetic lethal paralogs has existed for nearly a century^6^, the majority of paralogs remain uncharacterized and only recently have genomic sequences for multiple organisms enabled the prediction of synthetic lethal interactions^3^. Additionally, the number of potential genetic interactions across all paralog families is immense (>10^12^ interactions) and prohibitive for comprehensive genetic screening^4^. The expansion of genomewide loss-of-functions (LoF) CRISPR screens has uncovered novel synthetic lethal interactions, yet these data are limited by the incomplete representation of LoF events in human patients cancer cell models and therefore are likely to miss patient-relevant synthetic lethal interactions^2,7^. Alternatively, while synthetic lethal interactions of select candidate pairs have been tested through combinatorial LoF CRISPR screening, these pairs are typically limited to a small subset of the known human paralogs. As such, the potential functional interactions of most paralog pairs in the human genome remain unknown.

Here, we present the first genetic screen of the 36,648 known paralog pairs in the human genome using a multiplexed CRISPR-Cas12 library, which identified multiple synthetic lethal and buffering interactions. This unbiased screen also confirmed a wide variability in the penetrance of synthetic lethal interactions across different cell lineages and genetic backgrounds, suggesting that other endogenous factors influence the sensitivity to synthetic lethal mechanisms. We used XGBoost^8^ to characterize the predictive features of paralog synthetic lethality across three combinatorial CRISPR studies of 461 paralog pairs tested across 49 cancer cell lines. This integrated dataset represents the largest cohort of cancer models that have been screened for paralog synthetic lethalities and provided molecular insights to the cellular contexts that modify complex synthetic lethal interactions. We also demonstrate that the strength of synthetic lethal interactions are largely due to the essentiality of the protein-protein interaction networks shared by the paralogs pairs. Collectively, this report provides key molecular insights governing the penetrance and complexity of the synthetic lethal interactions of paralogous gene pairs.

## RESULTS

### Mapping genetic interactions of 36,648 paralog pairs in the human genome

The combined previous attempts to characterize paralog synthetic lethalities have been limited to testing approximately 20% of the human paralog pairs due to size restrictions of conventional multiplexed CRISPR libraries^9^. Moreover, roughly two-thirds of these paralog pairs were tested in only one cell model^9^, indicating that most of the human paralogome remains uncharacterized. To overcome these limitations, we developed a CRISPR-Cas12 screening approach to test all 36,648 paralog pairs in the human genome using a multiplexed library design with six independent constructs per paralog pair (**Table S1**). Paralog pairs were queried from Ensembl and required to have at least 20% homology, as described previously^10^. The CRISPR-Cas12 library design utilized arrays of four guides per construct and two guides per paralog, which performed optimally for combinatorial screens^9,11,12^ and enabled a minimal library size (n = 113,502 constructs) for screening all paralogs (n = 13,128 singletons and n = 36,648 pairs) (**Figure 1A**). Digenic screening of all human paralog pairs was performed using NCI-H1299 and MDA-MB-231 cells that stably expressed enAsCas12a, and synthetic lethal interactions were identified by comparing the observed effects of the double knockout (KO) with the additive effects of each single KO (**Figure 1B** and **Table S2**). Library performance was assessed by comparing the effects of gene KOs in the CRISPR-Cas12 library with the DepMap essentiality profiles of NCI-H1299 and MDA-MB-231. A receiver operating characteristic (ROC)–area under the curve (AUC) analysis revealed that the CRISPR-Cas12 performed well in detecting DepMap essential genes in MDAMB231 (AUC = 0.92) and NCI-H1299 (AUC = 0.90) (**Figure 1C**). Finally, a null-normalized mean difference (NNMD)^13^ analysis revealed slightly better separation of essential and non-essential genes in the CRISPR-Cas12 library relative to the DepMap (**Figure 1D**), potentially owing to the combined activity of the multi-guide constructs with enAsCas12a.

**Figure 1.**
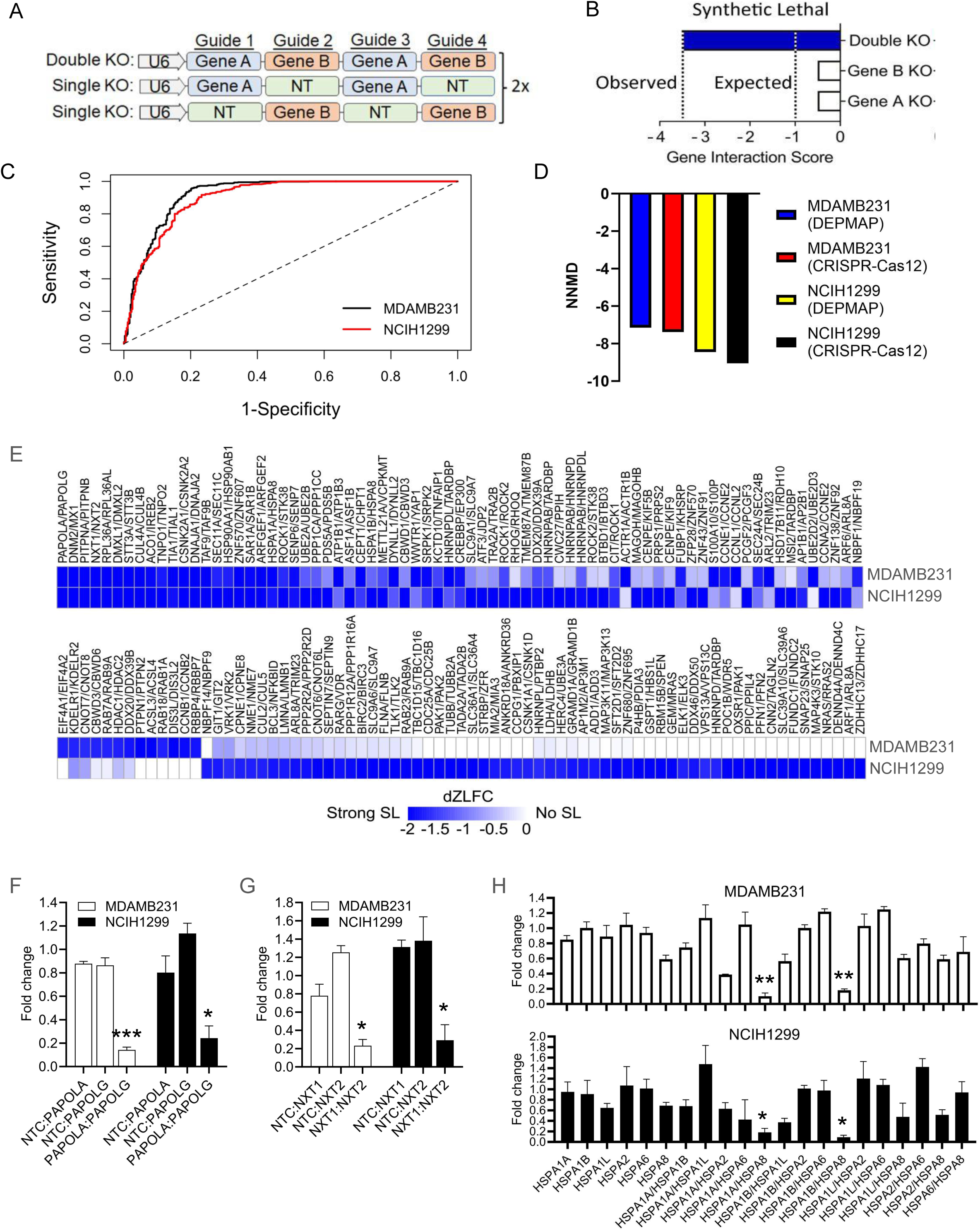
Multiplexed CRISPR/Cas12 screening of synthetic lethal interactions across all 36,648 paralog pairs in the human genome. (**A**) Schematic of the CRISPR/Cas12 library multiplexed guide arrays targeting one or two genes per array. (**B**) Schematic of the synthetic lethality screening approach using the CRISPR/Cas12 library. (**C**) Library performance was assessed by a receiver operating characteristic (ROC)–area under the curve (AUC) analysis to test the ability of the singleton constructs in the CRISPR/Cas12 library to predict gene essentiality as defined by the DepMap. (**D**) Library performance was also assessed by NNMD analysis of the delta of the top 5% essential genes (positive set) and bottom 5% non-essential genes (negative set) normalized to the standard deviation of the negative set. (**E**) Heatmap of dzL2FC values of the synthetic lethal interactions detected from 36,648 paralog pairs tested in NCI-H1299 and MDA-MB-231. (**F, G**) Examples of validated synthetic lethal interactions, PAPOLA/PAPOLG and NXT1/NXT2, respectively. \*\*\**P* < 0.001 and \**P* < 0.05, as determined by a 1-way ANOVA with Sidak’s post-hoc test. (**H**) An example of a previously unreported family (Hsp70) with a subset of synthetic lethal paralogs. \*\**P* < 0.01 and \**P* < 0.05, as determined by a 1-way ANOVA with Sidak’s post-hoc test.

A meta-analysis of five recent digenic paralog screens revealed that synthetic lethal interactions are highly heterogeneous and no synthetic lethal interactions were common across all tested cell models^9^. Moreover, out of the 6,868 paralog pairs that were tested, only 388 (5.6%) paralog pairs were identified as synthetic lethal in at least one cell mode^9^. Since these libraries were bioinformatically enriched for paralog subsets with the highest probability of synthetic lethal interactions, we sought to determine the overall prevalence of paralog synthetic lethalities amongst all known paralog pairs. Out of the 36,648 paralog pairs that were tested in NCI-H1299 and MDAMB231, surprisingly few synthetic lethal interactions (<0.4% of tested) were observed (144 and 45 pairs, respectively). In comparison, a markedly higher proportion of the 12,312 unique paralogous genes were essential singletons (∼3% of tested) in NCI-H1299 (n = 406 singletons) or MDA-MB-231 (n = 384 singletons). Of the 224 essential singletons that were shared by both cell lines, 193 were classified as common essential by the DepMap (86%). Collectively, these data suggest that pairwise paralog synthetic lethalities contributed to a relatively small proportion (∼10%) of the overall dependency profile of a cancer model.

Notably, of the 93 unique synthetic lethal paralog pairs that were detected in either cell model, 40 paralog pairs were novel and 53 have been replicated elsewhere (**Figure 1E** and **Table S3**), demonstrating the ability of the CRISPR/Cas12 library to detect pairwise synthetic lethal interactions. It was also notable that 17 out of the 22 (77%) synthetic lethalities that were detected in both cell models have been replicated elsewhere (**Table S3**), including PAPOLA/PAPOLG^14^ and NXT1/NXT2^15^ (**Figure 1F, G**). Likewise, the top ranking common synthetic lethalities (e.g., CSNK2A1/CSNK2A2, SAR1A/SAR1B, ARFGEF1/ARFGEF2, TIA1/TIAL1) overlapped with a gold standard set of synthetic lethal paralog pairs described elsewhere^9^. Multiple novel synthetic lethal interactions were also detected, including members of the heat shock protein family A (Hsp70) members, HSPA8/HSPA1A and HSPA8/HSPA1B, which can chaperon the same client proteins interchangeably^16–18^ (**Figure 1H**). Notably, the loss of other Hsp70 family members was not synthetic lethal (**Figure 1H**), suggesting that the combined loss of these paralogs was tolerable in these cellular contexts. Collectively, these data highlight the power of unbiased paralog mapping to discover synthetic lethal interactions, as well delineate lethalities within large families, such as the HSP70 family.

In addition to synthetic lethalities, combinatorial CRISPR screens also capture positive interactions^19^, whereby the loss of two paralogs results in a net positive advantage that is greater than the sum of either paralog alone (i.e., synthetic buffering). To test this possibility across all paralog pairs in the human genome, digenic buffering interactions were identified by comparing the observed effects of the double knockout (KO) with the additive effects of each single KO (**Figure 2A**). Synthetic buffering interactions of 18 paralog pairs were observed in NCI-H1299 (n = 17) or MDA-MDA-231 (n = 6), of which five buffering pairs were shared across both models (**Figure 2B**). Notably, 11 of 18 (61%) of the unique buffering pairs were closely associated by STRING analysis with tumor suppressor pathways, including MOB1B/MOB2 (**Figure 2C**, **D**), ARHGAP12/ARHGAP21 (**Figure 2E**, **F**), and EXOSC7/EXOSC8 (**Figure 2G**, **H**). Although these proteins, such as MOB1B and MOB2, have been individually associated with tumor suppressor pathways^20^, this is the first evidence of functional compensation by these tumor suppressor paralogs. For example, MOB1B and MOB2 are members of the MOB family (monopolar spindle-one-binder proteins) with distinct and overlapping roles in regulating the Hippo pathway^20^. Taken together, these data indicate the presence of functionally redundant paralog pairs, such as MOB1B and MOB2, exist within tumor suppressor pathways and likely convey a selective advantage for tumor cells upon loss of both paralogs^9^.

**Figure 2.**
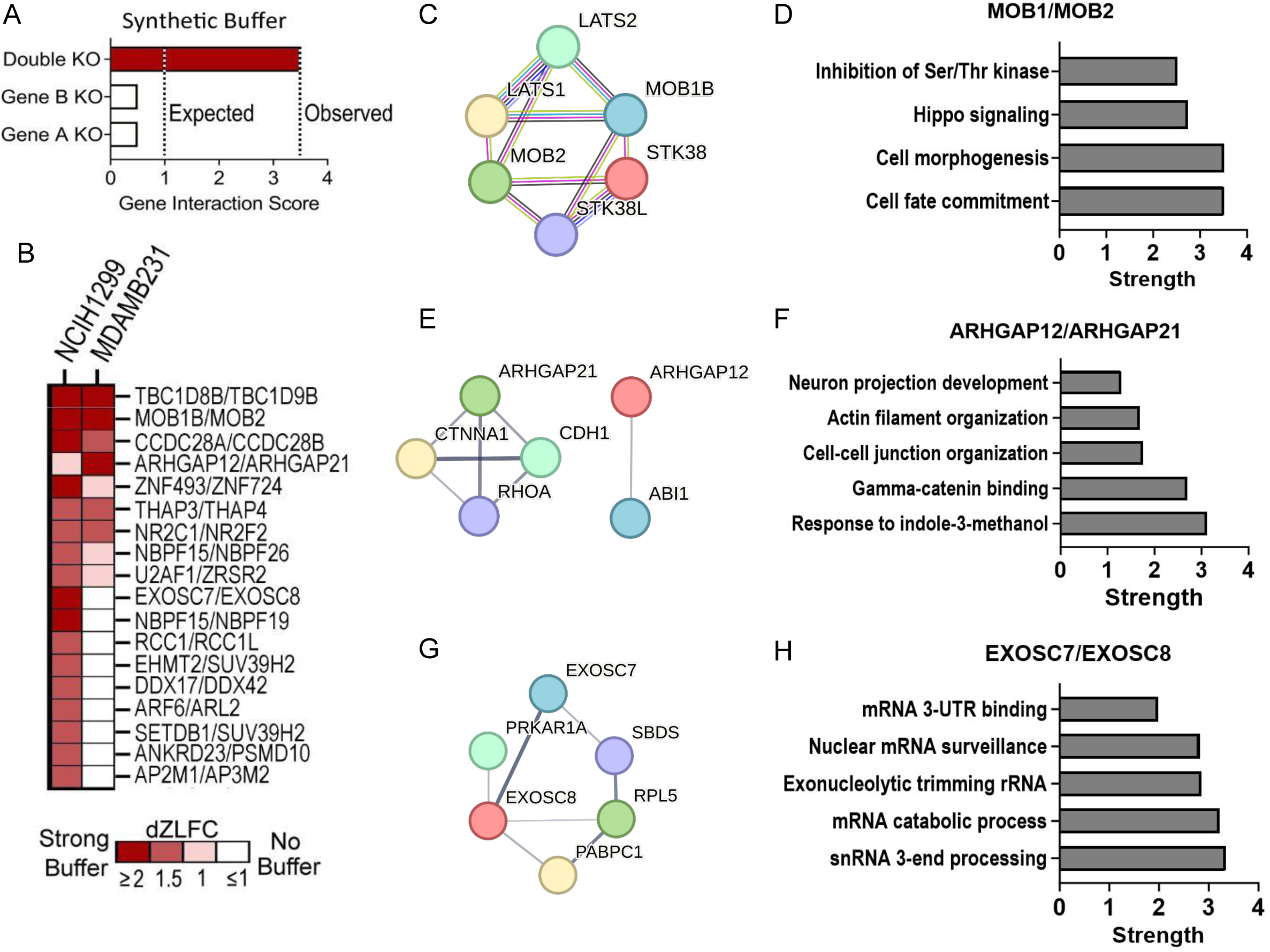
Multiplexed CRISPR/Cas12 screening of synthetic buffering interactions across all 36,648 paralog pairs in the human genome. (**A**) Schematic of the synthetic lethality screening approach using the CRISPR/Cas12 library. (**B**) A heatmap of synthetic buffering interactions in NCIH1299 and MDAMB231 cells. (**C**-**H**) STRING network and pathway enrichment analysis of synthetic buffering pairs, MOB1/MOB1B (MOB4) (**C**, **D**), ARHGAP12/ARHGAP21 (**E**, **F**), and EXOSC7/EXOSC8 (**G**, **H**). Note the connections with established TSGs and related pathways.

### Context dependency of paralog synthetic lethalities

Genetic mapping of all known human paralog pairs revealed relatively few synthetic lethalities that were common between cancer models, a phenomenon that has also been observed in more focused screening efforts^3,12,21^. One possible explanation is that most synthetic lethalities are complex polygenic interactions that are modified in different cellular contexts and by other endogenous factors^4^. We hypothesized that disentangling complex polygenic synthetic lethalities could be achieved by correlating pairwise synthetic lethalities with endogenous factors across many cell models. However, a barrier to this approach remains in the practical limitations of screening large multiplexed CRISPR libraries across many cell lines.

Since screening all paralog pairs across many cell lines would be technically challenging, we instead focused on screening the 5,116 paralog pairs with solved structures, as these are typically well-studied proteins with defined functional domains. To further streamline screening, a minimal CRISPR-Cas12 library was designed to interrogate dKO effects with four guides per construct and two guides per paralog (n = 10,886 constructs) (**Table S4)**. The average NNMD score across the 20 cell models (-4.1 ± 0.4) indicated that the library was sensitive in detecting a gold standard set of synthetic lethal paralog pairs^9^. The dKO effects of this compact library were then compared with the additive effects of each single KO in the Cancer Dependency Map (https://depmap.org/portal/), which enabled efficient screening across 20 cancer models. An ROC-AUC analysis revealed that synthetic lethal interactions could be accurately detected (AUC = 0.77) by comparing the normalized observed effects of the double KO (CRISPR-Cas12) with the normalized expected effects of the single KOs in the Cancer Dependency Map (**Figure 3A**).

**Figure 3.**
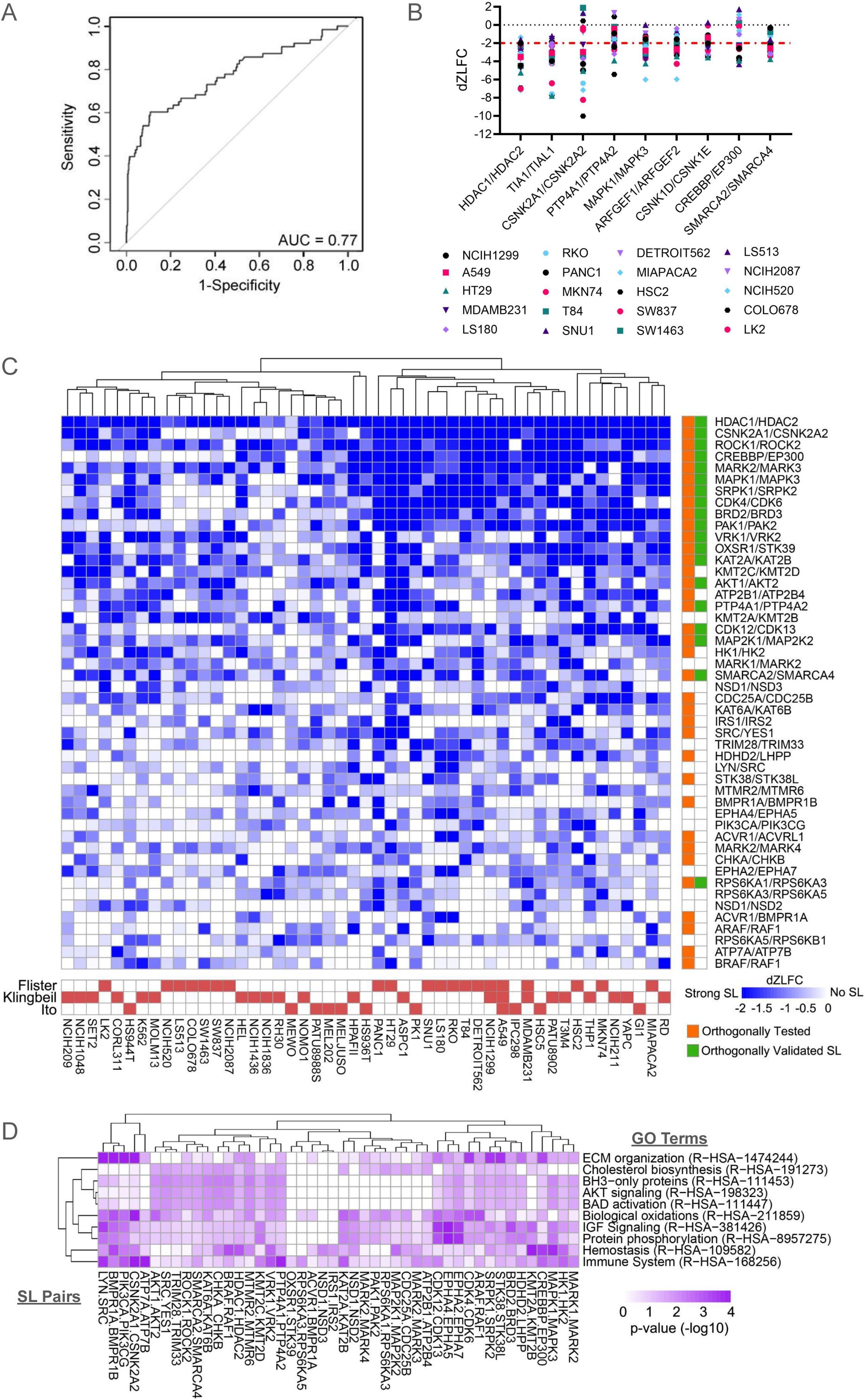
Meta-analysis of paralog synthetic lethalities across cancer models. (**A**) An ROC-AUC analysis revealed that synthetic lethal interactions could be accurately measured using a hybrid approach that compared the normalized observed doubled KO effects using multiplexed CRISPR-Cas12 with the normalized expected effects of the single KOs in the DepMap. (**B**) The hybrid screening approach detected all of the gold standard set controls that were included in the library. (**C**) A meta-analysis of synthetic lethal interactions that were detected in at least two of the 49 unique cancer models that were tested. (**D**) A heatmap of the p-values (-log10) of the most frequently enriched gene ontologies in the features selected by the XGBoost analysis to predict the synthetic lethal interactions depicted in (**C**). Pathway names corresponding to the HSA pathway numbers were abbreviated and the full pathway names are provided in Table S8.

A total of 780 synthetic lethal interactions were detected in at least one cell line out of 5,318 paralog pairs that were tested (14.6%), with wide variability in the number of synthetic lethalities per cell line (n = 17 - 230; median = 115) (**Table S5**). These synthetic lethal interactions could be classified as common (n = 63; >33% incidence), selective (n = 301; 33% to 10% incidence), and rare (n = 446; <10% incidence). The previously reported ‘gold standard’ synthetic lethal pairs^9^ that were included in this library were replicated as the most frequent synthetic lethal interactions (**Figure 3B**). For example, the histone deacetylases, HDAC1 and HDAC2, were the most commonly detected synthetic lethal interactions in 18 out of 20 cancer models (**Figure 3B**), agreeing with multiple other reports^14,21,22^. Likewise, 17 of the top 30 synthetic lethalities that were observed in ten or more cancer models have also been reported previously, including SMARCA2/SMARCA4^14,23^ and EP300/CBP^24^ (**Figure 3B**). This approach could also distinguish synthetic lethal interactions within family members, such as the synthetic lethal interactions of HSPA8 with HSPA1A or HSPA1B (**Supplemental Figure 1**), as was detected in NCIH1299 and MDAMB231 (**Figure 1E and 1H**). In contrast, the loss of HSPA8 was only synthetic lethal with the loss of HASPA1L, HSPA2, or HSPA6 in other cancer models (**Supplemental Figure 1**). Collectively, these data demonstrated that this hybrid approach was robust in detecting synthetic lethal interactions that are heterogenous in different cellular contexts.

Two similarly sized digenic paralog screens were recently published using combinatorial CRISPR-Cas9 libraries^21,25^, providing an opportunity to expand the cohort of cell lines for identifying the cellular contexts that modify synthetic lethal interactions. Following integration and normalization of the three studies, a total of 84 synthetic lethal interactions were detected out of 461 paralog pairs that overlapped between the three studies (**Table S6**). As reported previously^9^, no synthetic lethal interactions were completely penetrant, with HDAC1/HDAC2 remaining the most common synthetic lethality in 39 out of 49 cell lines tested (**Figure 3C**). The XGBoost machine learning method with L1 and L2 regularization was used to identify the cellular contexts that modify the strengths of digenic synthetic lethal interactions, including genomewide gene expression, essentiality, copy number variants, and somatic mutations, as input variables^8^. Despite including the four feature classes, only gene expression features were selected by the models (**Table S7**), fitting with previous observations that tumor dependencies are best predicted by transcriptomics data^12^. A gene ontology (GO) enrichment analysis was then performed using the selected features for the synthetic lethal paralog pairs detected in at least two cancer models (**Figure 3C** and **Table S8**). The GOs that were most frequently associated with synthetic lethal paralog pairs are shown in **Figure 3D**, including multiple key biological processes involved in oxidative stress and pro-survival pathways, such as AKT signaling and anti-apoptotic BH3 proteins. Notably, several GOs, such as extracellular matrix (ECM) and immune signaling, were associated with the strength of synthetic lethal interactions between paralog pairs that have been individual associated with these processes (e.g., LYN/SRC and BMP1RA/BMP1RB). Combined, these data demonstrated that endogenous cellular context modified the sensitivity to synthetic lethal interactions, which can be mapped across a panel of cancer models.

### Predicting paralog synthetic lethality

Although the majority of human genes are paralogous, only ∼20% have been empirically tested for synthetic lethal interactions prior to this study^9^. As such, computational approaches for predicting synthetic lethality have been limited by the lack of a robust and unbiased set of experimentally validated synthetic lethalities for training and evaluation of predictive models. To our knowledge, this study represents the largest set of paralog screens to date, which we hypothesized could be used to train an ensemble classifier of paralog synthetic lethality. To account for the variability in synthetic lethal interactions across different cellular contexts, weighted sets of true positives (n = 223 pairs per set) based on the overall representation across six distinct CRISPR screening datasets. These types of unbalanced positive-only learning tasks can be addressed by using randomly-sampled negative training sets of equivalent size to the positive set. Accordingly, size-matched negative sets were randomly sampled without replacement from the remaining set of paralog pairs without observed lethalities and a random forest model was built using the averaged predictions over 10,000 models that incorporated 25 sets of features. The feature sets spanned three broad categories (sequence, neighborhood, and expression) that were predefined to capture unique aspects of synthetic lethal biology^3^, as well as an additional fourth category that incorporated protein structural similarity from AlphaFold^26^. The resulting averaged model was then used to assign the probability of a synthetic lethal interaction between a paralog pair.

As shown in **Figure 4A-B**, the probability of synthetic lethal interactions were correlated with the prevalence of the observed synthetic lethality, demonstrating that the random forest model is able to differentiate common, selective and rare synthetic lethal interactions. The average decrease in the Gini metric was used to assess the relative importance of the features in the models, which revealed that the extent of protein-protein interactions (PPI) of the paralog pairs and the essentiality of the interaction network were strong predictors of synthetic lethality (**Figure 4C**). The average expression of either paralog in normal tissues was also highly predictive of paralog synthetic lethality (**Figure 4C**). The SHapley Additive exPlanations (SHAP) scores were also calculated to determine the relative contribution of the features to individual paralog pairs (**Table S9**), which confirmed the importance of PPI networks and normal tissue expression in predicting synthetic lethality (**Figure 4D-F**). Moreover, comparing the SHAP scores across the classifications of common, selective, and rare synthetic lethal interactions revealed that the size and essentiality of the network of PPI shared by the paralog pairs made the largest contributions to whether a synthetic lethality was common across cancer models (**Figure 4G**, **H**). To test this hypothesis further, the average essentialities of the 780 synthetic lethal interactions that were classified as common (n = 63; >33% incidence), selective (n = 301; 33% to 10% incidence), and rare (n = 446; <10% incidence), based on experimental observations across the 20 cancer models. This revealed that the shared PPI network of common synthetic lethal paralog pairs had significantly more essential genes on average (n = 19 per pair) than selective synthetic lethalities (n = 6 per pair) or rarely synthetic lethal pairs (n = 3 per pair) (**Figure 4I**). Several known broadly acting synthetic lethal paralog pairs were ranked in the top most essential PPI networks among common synthetic lethalities (**Figure 4J**), as were multiple synthetic lethal paralog pairs in Hsp70 family, including HSPA2/HSPA8 (**Figure 4K**). Taken together, these data demonstrated that the essentiality of a PPI network that is shared by a paralog pair is a major determinant of the commonality of the synthetic lethality.

**Figure 4.**
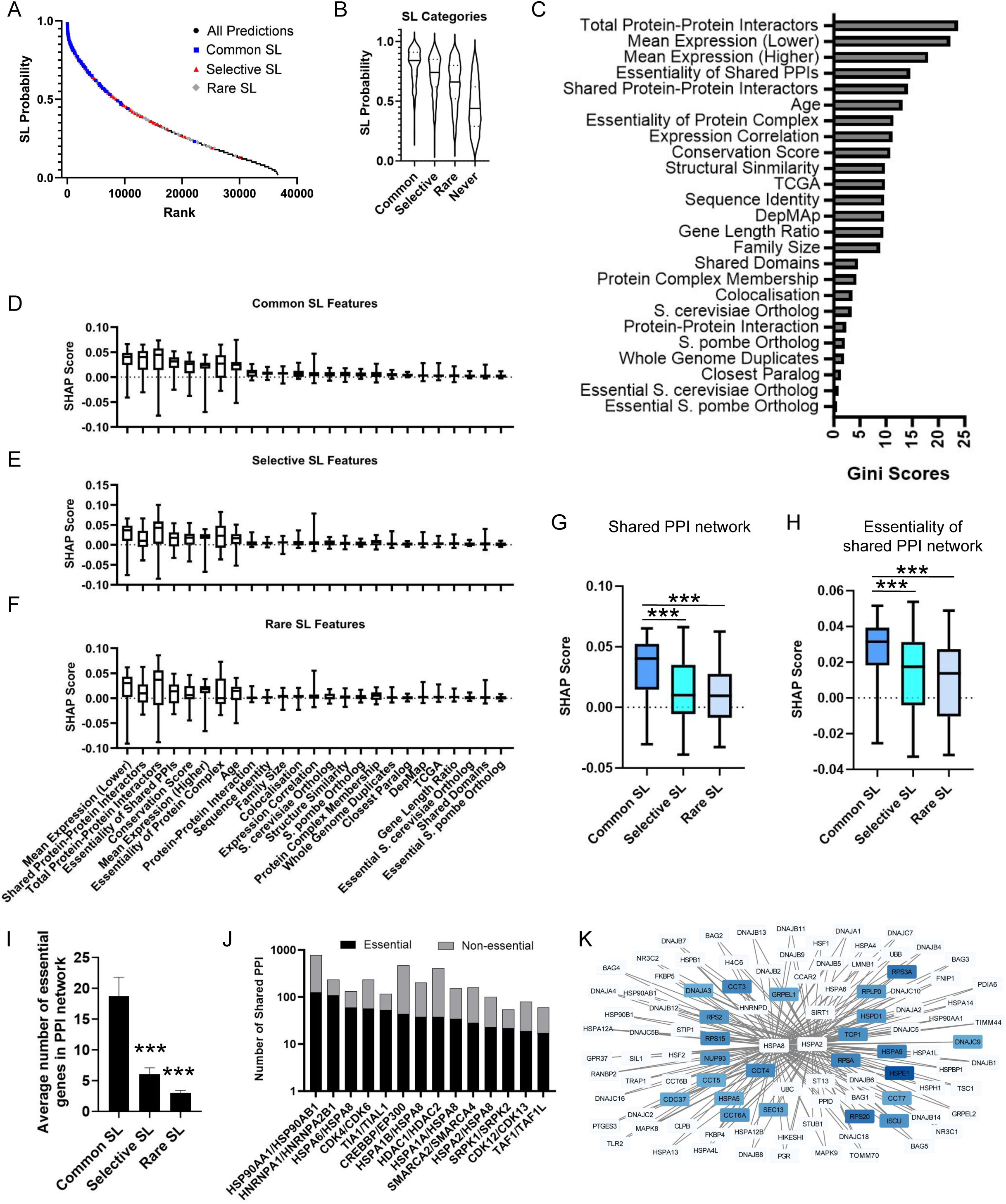
A weighted random forest classifier for predicting digenic synthetic lethal paralogs. (**A**) The distribution of synthetic lethality probability scores across the 36,648 paralog pairs in the human genome. (**B**) A violin plot showing the distribution of probability scores across synthetic lethal paralog pairs that were classified as common (>33% prevalence), selective (33%-10% prevalence), rare (<10% prevalence), or never detected. (**C**) The mean decrease in the Gini metric was used to assess the relative importance of the features in the model. (**D**-**F**) The distribution of SHAP scores across paralog synthetic lethalities that were classified as common, selective, or rare. (**G**, **H**) The comparison of SHAP scores across classes revealed that the size (**G**) and essentiality (**H**) of the PPI network shared by a paralog pair are the major contributors to whether a synthetic lethality was common across cancer models. \*\*\**P* < 0.001, as determined by one-way ANOVA with Tukey’s multiple comparison test. (**I**) Average number of essential genes in the shared PPI network of synthetic lethal interactions classified as common (n = 63; >33% incidence), selective (n = 301; 33% to 10% incidence), and rare (n = 446; <10% incidence). Data are presented as mean +/- SEM. \*\*\**P* < 0.001, as determined by one-way ANOVA with Dunnett’s multiple comparison test. (**J**) The distribution of essential and non-essential genes in the shared PPI network of the top common synthetic lethal paralog pairs. (**K**) Plot of the PPI network shared by the synthetic lethal paralog pairs, HSPA2 and HSPA8. Essential genes (DepMap Chronos Score < -1) are indicated by blue shading.

## DISCUSSION

This study addressed the significant challenge of empirically testing the genetic interactions of all known paralog pairs in the human genome, which has been limited by the experimental capacity of existing technologies for combinatorial genetic perturbation screens^4^. To overcome these limitations, we developed a CRISPR-Cas12 screening approach, which performed well in detecting synthetic lethal and buffering interactions in two distinct cancer models. This study revealed that synthetic lethal interactions are relatively rare on a per tumor basis (∼0.4% of all paralogs). In addition to rarity, a meta-analysis of a subset of 461 paralog pairs in 49 cancer models confirmed that the penetrance of synthetic lethal interactions is variable, and in some cases, can be predicted by polygenic interactions with endogenous factors. Acknowledging that the ability to predict digenic interactions is likely limited by the variability across cancer models, we built a machine learning classifier that utilized a true positive set that was weighted by the penetrance of synthetic lethal interactions across cancer models. The weighted machine learning classifier revealed that the penetrance of a synthetic lethal pair is modified by the overlap and essentiality of the mutual PPI network that interacts with a paralog pair, which is both a novel and intuitive finding. Thus, taken together, this study provides a comprehensive analysis of all digenic interactions in the human genome, as well as the key features that underlie the heterogeneity in synthetic lethalities that have been reported here and elsewhere.

### Pairwise interactions of paralogs are heterogenous and relatively rare

Multiple studies have demonstrated a wide variability in the penetrance of digenic synthetic lethal interactions using focused CRISPR screens^12,14,19,21,27,28^, yet interpreting these data have been difficult due to the limited overlap in the paralog sets and technologies used in each study^29^. Here, we screened all 36,648 paralog pairs in the human genome using a single gene-editing technology (enAsCas12a), which confirmed unambiguously that a minority of synthetic lethalities (∼10%) are shared by two cancer models. Fitting with recent observations that few synthetic lethal interactions are common in all contexts^29^, the expanded meta-analysis of 49 cancer models revealed no synthetic lethalities that were completely penetrant, with HDAC1/2 reaching the highest penetrance of 80%. We hypothesized that the penetrance of synthetic lethal interactions was modified by cellular context, which was supported by a XGBoost analysis that identified the predictive features of some digenic synthetic lethalities. Notably, the most predictive features typically fell within a related cellular function, suggesting that these synthetic lethalities are likely complex polygenic interactions that are context dependent. These data agree with several reports of digenic synthetic lethality that were dependent on driver mutations within the pathway^21,30^, as well as the recently reported IN4MER approach that identified several trigenic synthetic lethalities^29^. Taken together, the data presented in our study confirmed and extended the examples of polygenic synthetic lethalities through complex interactions with endogenous factors in different cancer models.

Another key finding of this study was that digenic paralog synthetic lethalities are relatively rare when compared to monogenic dependencies. Approximately 0.4% of all 36,648 paralog pairs in the human genome were synthetic lethal in MDAMB231 or NCIH1299 cells, whereas ∼3% of the tested singletons were essential in either model. Even fewer paralog pairs were synthetic lethal in both cancer models (0.06%), while the majority of essential singletons were shared by both models, and were defined as “common essential” by the DepMap. To our knowledge, the meta-analysis of the 49 cancer models represents the largest combined screen of a subset of paralog pairs to date, providing an opportunity to more broadly assess the penetrance of synthetic lethality across different cellular contexts. Notably, this meta-analysis confirmed that the vast majority of digenic synthetic lethalities were rarely detected in multiple cancer models, which could be explained by complex polygenic interactions that vary based on cellular context and endogenous features^4^. Alternatively, these observations might also suggest tumors evolve specific programs of paralog synthetic lethalities based on lineage and genetic drivers, as has been proposed recently^10^.

In addition to synthetic lethal interactions, the 36,648 paralog screen also captured positive interactions (i.e., synthetic buffering), whereby the loss of two paralogs drives a net positive advantage in cancer cells. Similar to a previous report^19^, we found that the synthetic buffering paralog pairs were extremely rare when compared to synthetic lethal interactions and overlapped strongly with tumor suppressor genes (TSGs). Combined with the observation that paralogs are more likely to be homozygously deleted with TSGs^10^, these data suggest that synthetic buffering by paralogs is likely a mechanism that enables tumorigenicity. Moreover, based on the frequency of homozygous deletions in TCGA patients^31^, it is likely that the rarity of synthetic buffering is driven by the incidence of homozygous deletions of TSGs.

### Predicting digenic synthetic lethality of paralogs

Multiple attempts have been made to predict paralog synthetic lethality^3^, yet these efforts have been limited by an incomplete set of validated synthetic lethal interactions for the training and evaluation of machine learning models. De Kegel et al^3^ recently reported what was arguably the most comprehensive study of paralog synthetic lethality to date, prompting us to adopt a similar approach with several key changes. Firstly, rather than focusing the training sets (both positive and negative) on observations made from genetic interactions within the DepMap^3^, the positive training set used in our approach was defined by empirical observations and the negative set was represented as a random sampling of all other paralog pairs. Thus, the machine learning classifier used in the present study is distinct in that the training sets were not reliant on a limited representation of synthetic lethalities in the DepMap^7^. Secondly, our machine learning classifier was unique in that it used a weighted positive-only training set, which accounts for the widely variable penetrance of paralog synthetic lethalities that was reported here and elsewhere^12,14,19,21,27,28^. To our knowledge, this new classifier revealed for the first time that the overlap and essentiality of the PPI network that is shared by a paralog pair are the best predictors of the most penetrant synthetic lethalities.

### Moving beyond digenic synthetic lethal interactions

The incomplete penetrance of synthetic lethal interactions suggests that most synthetic lethal interactions are not digenic and will require further exploration to understand the molecular underpinnings of synthetic lethality^4^. One approach used in this study was to test the association of a subset of 461 paralog pairs across 49 cancer models. Notably, this approach revealed multiple digenic synthetic lethalities that were associated with endogenous factors, suggesting the possibility that larger screens of digenic interactions will unveil additional endogenous modifiers of polygenic synthetic lethalities in the future. An empirical approach to testing and validating polygenic synthetic lethalities has recently been demonstrated using multiplexed CRISPR-Cas12, in which families of up to four paralogs could be empirically tested^29^. However, although this exciting new approach offers to expand the repertoire of polygenic synthetic lethal interactions, there remains practical limitations in the feasibility of testing all polygenic interactions^4^. Thus, to expand beyond digenic synthetic lethalities, it is likely that a combination of approaches will be required. These include the observation of polygenic interactions from digenic screens and empirical validation using multiplexed screening approaches to test two or more paralogs simultaneously.

### Conclusions

The concept of synthetic lethal paralogs has existed for nearly a century^6^, yet most human paralogs had remained untested. Here, we presented a screening approach to test synthetic lethalities across all 36,648 paralog pairs in the human genome. Combined with focused paralog synthetic lethality studies across 49 cancer models, this work identified >800 synthetic lethalities with varied penetrance across different cancer models, which in some cases could be related to endogenous perturbations of related pathways and biological processes. Based on the observation that the penetrance of digenic synthetic lethal paralogs varies widely and are likely polygenic in most cases (i.e., three or more interactions), we built a new machine learning classifier that weighted the most penetrant synthetic lethal paralog pairs. Thus, this new classifier is better suited to modeling the predictive features of broadly essential paralogs pairs, which subsequently revealed that highly penetrant digenic synthetic lethal interactions were predicted by the essentiality of the mutual PPI network of the paralog pair. This observation indicates that machine learning approaches could be further refined to predict polygenic synthetic lethalities. To this end, future work is needed to expand the catalog of polygenic synthetic lethalities beyond the few examples that have been reported in the literature^4^.

### Limitations of the study

While the presented data provided key insights to the molecular underpinnings of paralog synthetic lethality, several questions remain unanswered due to the current limitations. The variable strength of synthetic lethalities that was reported here and elsewhere^9^, suggests that additional screens across diverse cell models will be needed to comprehensively predict complex polygenic interactions (i.e., three or more interactions). We demonstrated the feasibility of this approach using a meta-analysis of a subset of paralog pairs tested across 49 cell models. Nonetheless, a comprehensive screen of all 36,648 paralog pairs across many cancer cell backgrounds is needed to characterize the complex polygenic interactions that likely exist. The outputs of these screens could then be used to identify paralog families of three or more interactions for empirical testing using a multiplexed CRISPR-Cas12 library designed to test up to four interacting genes per construct, as has been recently demonstrated elsewhere^9^. However, these libraries would be restricted in size to testing only portions of the human paralogome and would therefore require observation-based predictive modeling to select the most likely candidates to test *a priori*. As foundational datasets grow, further innovations in machine learning approaches will also be needed to predict potential synthetic lethal interactions. Finally, a major limitation of all paralog screens to date is the exclusive focus on essential functions, whereas it remains possible that the majority of synthetic interactions of paralogs impact other biological functions.

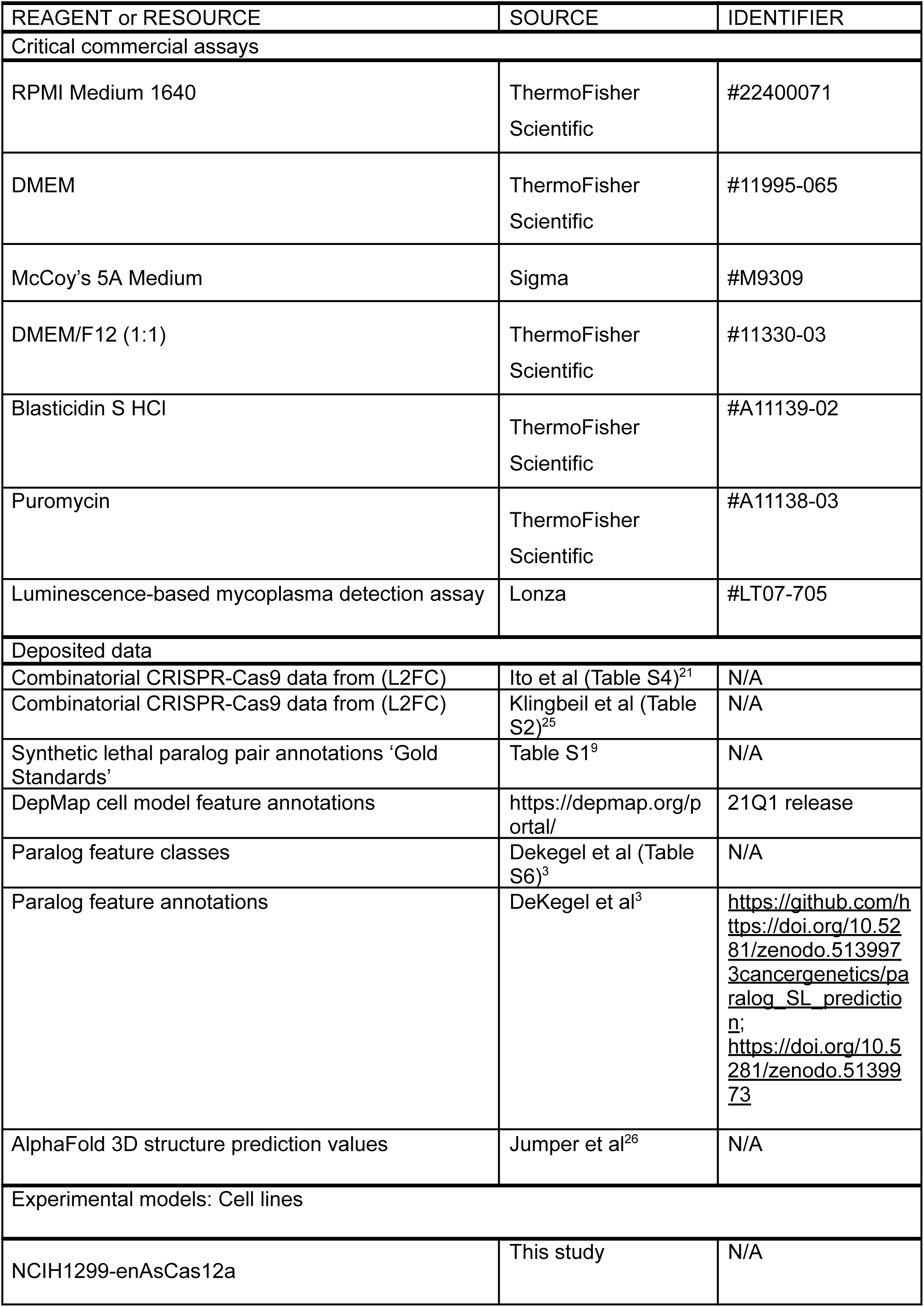
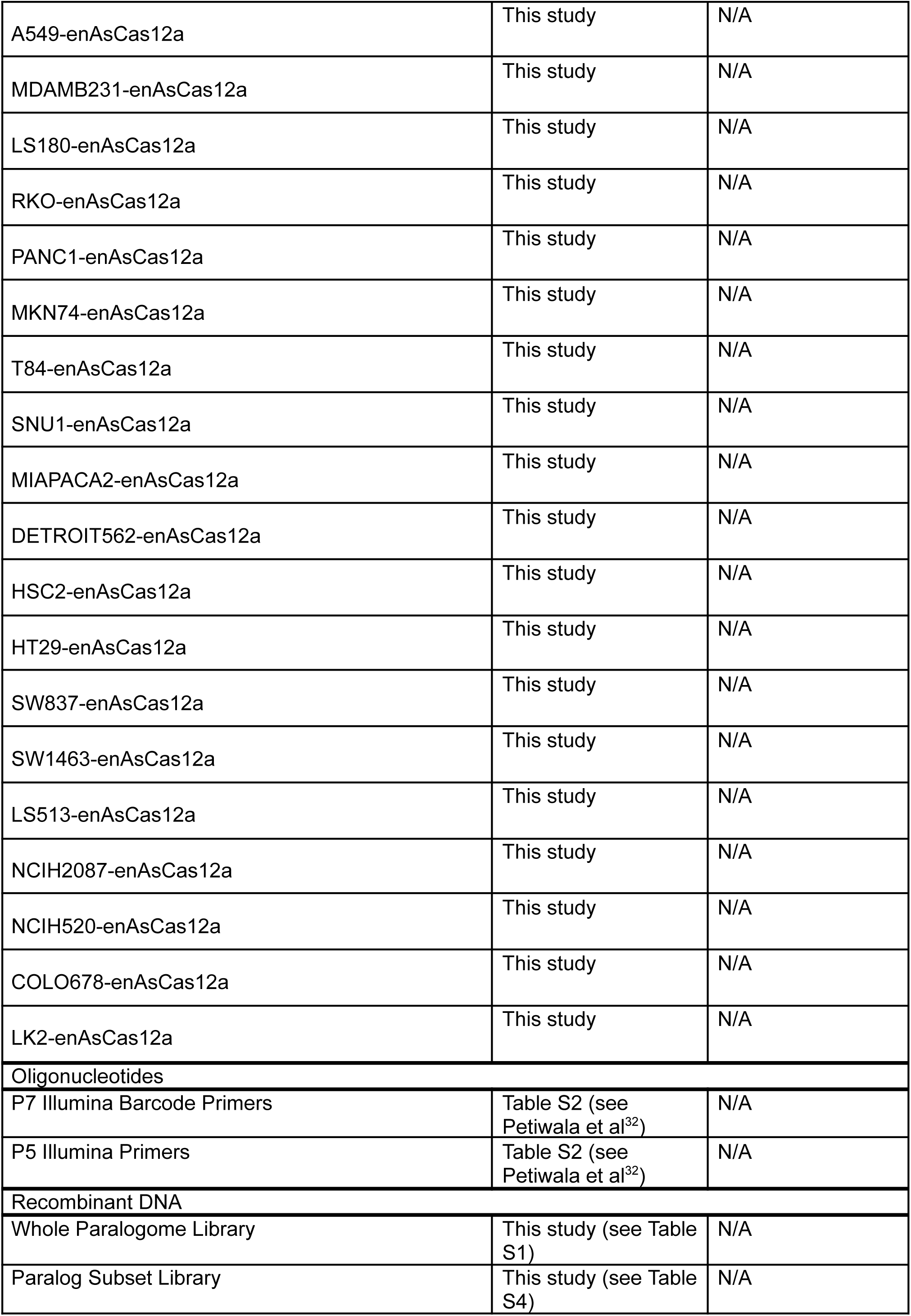
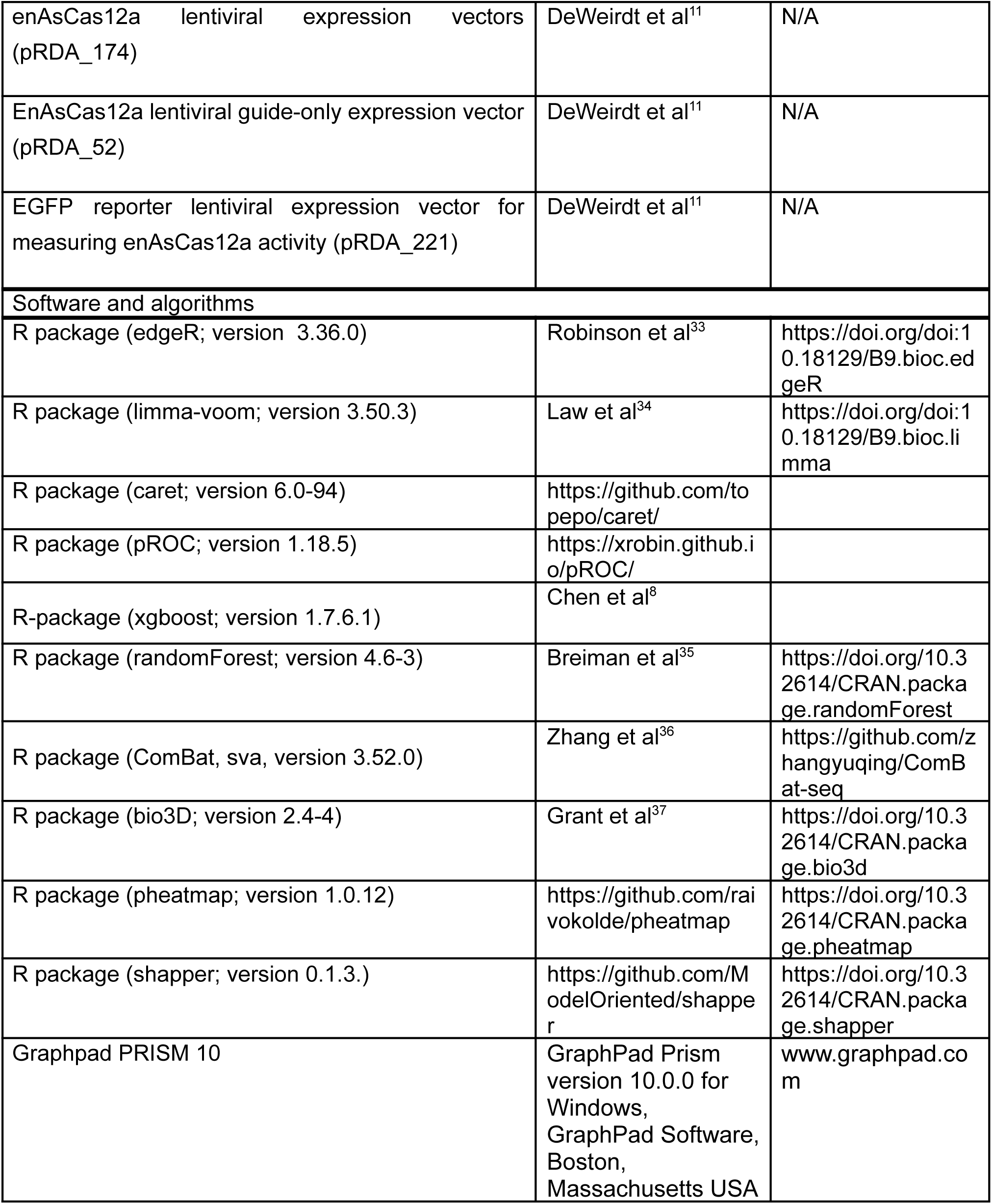

## EXPERIMENTAL MODEL DETAILS

### Cell Culture

The following cancer models were transduced with lentivirus stably expressing enAsCas12a with a blasticidin selection vector (pRDA_174): NCIH1299, A549, MDAMB231, LS180, RKO, PANC1, MKN74, T84, SNU1, MIAPACA2, DETROIT562, HSC2, HT29, SW837, SW1463, LS513, NCIH2087, NCIH520, COLO678, and LK2. Briefly, depending on the recommended media, cells were cultured in RPMI Medium 1640 (ThermoFisher Scientific, #22400071), DMEM (ThermoFisher Scientific, #11995-065), McCoy’s 5A Medium (Sigma, #M9309), or DMEM/F12 (1:1) (ThermoFisher Scientific, #11330-03) supplemented with 10% fetal bovine serum (FBS, ThermoFisher Scientific # 10082147) and 100U/ml penicillin/streptomycin (ThermoFisher Scientific #15140122). Following lentiviral transductions, cells were cultured in growth media with blasticidin (5ug/ml) (ThermoFisher Scientific, #A11139-02 and maintained in exponential phase growth by routine passaging in a 37 °C humidity-controlled incubator with 5.0% carbon dioxide. Routine testing for Mycoplasma was performed using a luminescence-based mycoplasma detection assay (Lonza #LT07-705).

### EGFP disruption assay to measure enAsCas12a activity in cell models

As described in previously^32^, parental and enAsCas12a expressing cells were infected with pRDA_221 lentivirus respectively that carries EGFP reporter sequence and a sgRNA targeting EGFP by spinfection in 12-well plates at 1000xg for 2hrs in presence of 10ug/ml polybrene at an MOI of 0.3. After an overnight incubation at 37°C, cells were selected for 3 days in puromycin (2 ug/ml). After selection, cells were cultured in presence of puromycin for another 3-4 days and then analyzed for EGFP expression using flow cytometry. All transgenic enAsCas12a models achieved >75% activity (**Table S10**).

## METHOD DETAILS

### Multiplexed pairwise screening of the human paralogome

Guides were designed for the enAsCas12a enzyme using the CRISPick (https://portals.broadinstitute.org/gppx/crispick/public) and synthesized into 4-guide arrays with direct repeats (DR)-0, -1, -2, and -3 preceding each guide, followed by cloning into a guide-only lentiviral vector (pRDA_052), as described previously^11,32^. Two double knockout constructs consisting of 2 guides x 2 genes (n = 4 guides total per construct) were designed for each pair of paralogs. Two single knockout constructs were also designed for each paralog pair: 2 guides x 1 gene + 2 non-targeting guides (n= 4 guides total per construct). A total of 500 constructs with 4 non-targeting guides were also included in the library as negative controls. The final library to test all paralog pairs in the human genome consisted of 113,502 constructs (n = 13,128 singletons and n = 36,648 pairs). Screens were performed in triplicate using NCI-H1299 and MDA-MB-231 cells that stably expressed enAsCas12a, as described previously^11^. Cells were infected at a multiplicity of infection (MOI) = 0.3 and cultured for 14 days while continuously maintaining 500X coverage, followed by DNA extraction and PCR-barcoding using the p5 Agon and p7 Kermit primers^11^. The PCR-barcoded libraries were single-end sequenced using an Illumina HiSeq4000 (300X cycle), followed by demultiplexing of sequencing reads (bcl2fastq, Illumina) and quantification of guide array abundance across all samples was done with a custom Perl script. Sequences between the flanking sequences or by location were extracted and compared to a database of sgRNA for each library. Only perfectly matched sgRNA sequences were kept and used in the generation of count matrix.

### Hybrid screening approach for testing a subset human paralogs

To test a subset of 5,116 paralog pairs by dKO, a CRISPR-Cas12 library design utilized arrays of four guides per construct and two guides per paralog (n = 10,886 constructs). Two double knockout constructs consisting of 2 guides x 2 genes (n = 4 guides total per construct) were designed for each pair of paralogs. Library synthesis, cloning, and screening was performed as described above using the following enAsCas12a transgenic cancer models: NCIH1299, A549, MDAMB231, LS180, RKO, PANC1, MKN74, T84, SNU1, MIAPACA2, DETROIT562, HSC2, HT29, SW837, SW1463, LS513, NCIH2087, NCIH520, COLO678, and LK2. Cells were infected at a multiplicity of infection (MOI) = 0.3 and cultured for 14 days while continuously maintaining 500X coverage, followed by DNA extraction, PCR-barcoding, and sequencing, as described above. Sequences between the flanking sequences or by location were extracted and compared to a database of sgRNA for each library. As described above, perfectly matched sgRNA sequences were used in the generation of the count matrix.

## QUANTIFICATION AND STATISTICAL ANALYSIS

### Analysis of multiplexed pairwise screening of the human paralogome

Normalization of the sgRNA count matrix between all samples was performed using the “TMM” method^38^ implemented in the edgeR R Bioconductor package^33^. Log2 fold-changes (L2FC) of guide array abundance were calculated by comparing day 14 libraries with the plasmid library using limma-voom^34^. Genetic interactions (GI) were calculated by comparing the expected and observed L2FC of double and single knockout constructs following z-score transformation (zL2FC)^12^. Paralog pairs were filtered for expression (>2.5 TPM) and an observed activity of the median double knockout effect of <-1 zL2FC and >1 zL2FC for synthetic lethal and synthetic buffering pairs, respectively. A paralog pair with a z-transformed delta zL2FC (dzL2FC) of >1.64 between the observed and predicted effects was considered significant. Screen performance was assessed using multiple methods, including the null-normalized mean deviation (NNMD) method, as described previously^13^. Briefly, the difference between the delta of the mean effects of the aggregate top 5% most essential genes (positive set) and the aggregate of the bottom 5% least essential genes (negative set) of the MDAMB231 and NCIH1299 cell models were normalized to the standard deviation of the negative set. Screen performance was also evaluated by ROC-AUC analysis to test the ability of the singleton constructs in the CRISPR/Cas12 library to predict gene essentiality as defined by the DepMap. A 10-fold cross-validation was used to train the generalized linear model classifier and predict the essentiality on the hold-out data for each cell line separately. The cross-validation was performed with the R package caret 6.0-94, using the “glm” method and “binomial” family option. The R pROC package (version 1.18.5) was used to plot the ROC curve and compute the AUC value.

### Analysis of hybrid screening approach for testing a subset of human paralogs

Normalization of the sgRNA count matrix between all samples was done using the “TMM” method^38^ implemented in the edgeR R Bioconductor package^33^ and the L2FC of guide array abundance were calculated by comparing day 14 libraries with the plasmid library using limma-voom^34^. To estimate single gene knockout effects, sgRNA count matrices for each cancer model were directly downloaded from the DepMap (AvanaRawReadcounts.csv) and L2FC changes of single gene effects were calculated as described above. The GI were calculated by comparing the expected and observed L2FC of double and single knockout constructs following z-score transformations. A paralog pair with a dzL2FC > 2 (95% confidence interval) between the observed and predicted effects was considered significant. A receiver operating characteristic (ROC) curve was used to assess the efficacy of the hybrid screening approach using CRISPR-Cas12 to measure double KO effects and the DepMap to measure single gene KO effects. The R pROC package (version 1.18.5) was used to plot the ROC curve and compute the AUC value. Screen performance was assessed by the NNMD method, as the delta of the mean effects of the gold standard set of synthetic lethal paralog pairs included in this study (HDAC1/HDAC2, TIA1/TIAL1, CSNK2A1/CSNK2A2, CSNK1D/CSNK1E, MAPK1/MAPK3, PTP4A1/PTP4A2, and ARFGEF1/ARFGEF2)^9^, and a negative set comprised of library constructs targeting intergenic sequences.

### Meta-analysis of 461 paralog pairs in 49 cancer models

The L2FC data were directly downloaded from Ito et al^21^and Klingbeil et al^25^, followed by z-transformation to yield zL2FC values. The GI scores were calculated, as described above, by comparing the expected and observed L2FC of double and single knockout constructs following z-score transformations. A paralog pair with a z-transformed dzL2FC > 2 (95% confidence interval) between the observed and predicted effects was considered significant. Following dzL2FC calculations, a principal components analysis (PCA) revealed that batch effects persisted between the three studies, potentially owing to the distinct library designs and the use of different CAS enzymes^21,25^. To remove batch effects, the dzL2FC data were post-processed using ComBat^36^. The gene expression, mutations, copy number variants, and essentiality data were downloaded for each available cancer model (n = 47) from the DepMap (release 21Q1). The gene expression data were filtered for a variance of greater than 0.5 across the cancer models (n = 11,412 genes). The mutation data were filtered for a frequency of greater than 2% across the cancer models (n = 9,157 mutations). The copy number variant data were filtered to identify deletions with a relative ploidy of less than 1.2 and a frequency of greater than 5% (n = 6,757 deletions). The xgboost library (R package 1.7.6.1)^8^ for feature selection on SLs that had at least one essential cell line. In applying the xgboost algorithm, we opted for the ‘gblinear’ booster with ‘squarederror’ objectives. L1 and L2 regularizations were incorporated to reduce the redundancy of the selected feature set. We employed the xgb.cv function for 5-fold cross-validation to determine the optimal alpha and lambda parameters, exploring values between 0 and 1 in 0.1 increments. The best-performing alpha and lambda values were then used in the final model training step. During training, we determined the optimal number of iterations using <u>xgb.cv</u>, starting with 100 rounds and utilizing early stopping after 5 rounds if no improvement was observed. The optimal alpha, lambda, and number of rounds were applied in the xgb.train function to construct the final model. Finally, the feature importance was extracted from the model using the xgb.importance function.

### Machine Learning Predictions of Synthetic Lethal Paralogs

The random forest models and predictions were performed using the *randomForest* (v4.6-3)^35^ and *caret* (v6.0-93) (https://github.com/topepo/caret/) libraries in R statistical software. The number of trees in each model was fixed at 1000, as any increase in trees above this value did not yield any significant changes in the prediction probability. The tuneRF function in the random forest package was used to determine the optimal number of variables (“mtry”) sampled at each split. The stepfactor was set at 0.01 and the “improve” parameter at 0.01 value. A weighted positive set was randomly sampled without replacement (n = 223 per set) to match the penetrance of the 825 synthetic lethal paralogs observed in this study. The weight was defined as the number of cell lines where the synthetic lethality was observed divided by the total number of cell lines tested. To balance the training set for the model, size-matched negative training sets were generated by random sampling without replacement of genes and excluding the positive training set. A total of 10,000 random forest models using each of the random negative sets were trained and predictions were made based on each model, followed by averaging the predictions over 10,000 models to assign the synthetic lethality score to each paralog pair. There were 25 features which included 22 from the DeKegel et al. study and three additional features. Two features were the correlation values of the paralog pairs based on the DepMap CRISPR screen values and the co-expression from the TCGA. An additional feature was included by calculating the structure similarity of the pairs based on the AlphaFold 3D structure prediction values^26^. The structure similarity was determined based on the root-mean-square deviation of atomic positions (RMSD) of the superimposed protein structures. The RMSD was calculated using the bio3D (2.4-4) R package. For feature importance both the Gini metric and the SHAP values were calculated. The Gini values were calculated using the R randomForest packaged and averaged over the 10,000 model to determine the final contribution. The SHAP scores were generated using the shapper R package. The contributions of features to each individual paralog pair were averaged over 1,000 models.

## DECLARATIONS

### Author Contributions

M.J.F., Z.D., C.G., and D.V. conceptualized the study. M.J.F, D.V., and S.P. performed the experiments. M.J.F., Z.D., X.S., C.G., D.V., S.P., and C.L. analyzed and interpreted data. M.J.F. and Z.D. wrote the original manuscript draft. All authors reviewed and edited the manuscript draft.

## Acknowledgements

We thank AbbVie employees Aparna Vasanthakumar, Archana Iver, Kari Barlan, and Kevin Zhao for the helpful discussions and critical feedback. We also thank Erin Murphy, Tifani Anton, and the other members of the Genome Technology team in the AbbVie Genome Research Center for their expertise and technical support.

