## Supplemental Figure 1 for "Genetic Mapping of All Human Paralog Pairs to Characterize Synthetic Lethality and Buffering"

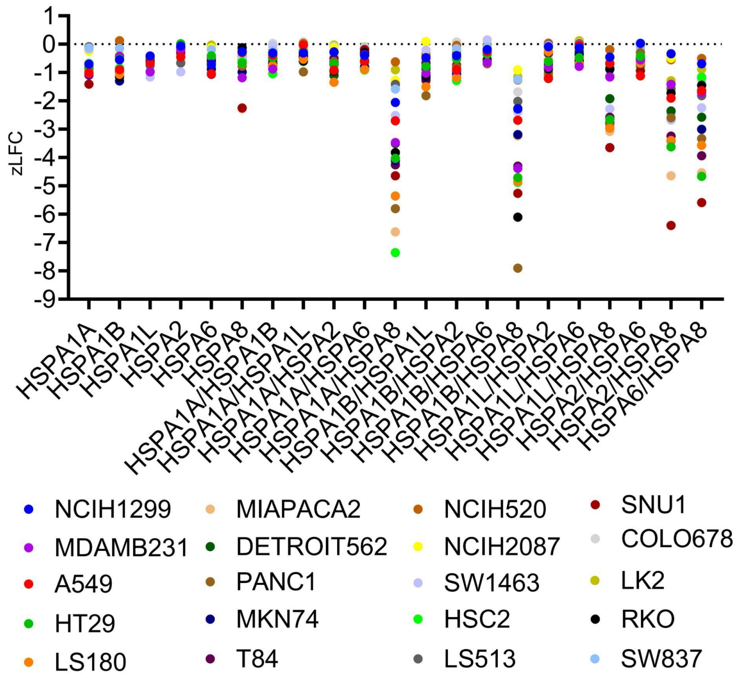


**Figure S1. The z-transformed log2 fold-changes of the HSP70 family members from the hybrid CRISPR-Cas12 screen of paralog pairs compared with the DepMap single-gene essentiality scores across 20 cancer cell lines. These data confirmed the synthetic lethal interaction of HSPA8 with HSPA1A and HSPA1B in MDAMB231 and NCIH1299 cells, as well as identifying additional synthetic lethal interactions with HSPA2 and HSPA6.**
